# Calcitic prisms of the giant seashell *Pinna nobilis* form light guide arrays

**DOI:** 10.1101/2022.10.25.513698

**Authors:** Shahrouz Amini, Tingting Zhu, Abin Biswas, Mohammad A. Charsooghi, Kyoohyun Kim, Simone Reber, Yannicke Dauphin, Peter Fratzl

## Abstract

The shells of the *Pinnidae* family are based on a double layer of single-crystal-like calcitic prisms and inner aragonitic nacre, a structure known for its outstanding mechanical performance. However, on the posterior side, shells are missing the nacreous layer, which raises the question of whether there could be any functional role in giving up this mechanical performance. Here, we demonstrate that the prismatic part of the *Pinna nobilis* shell exhibits unusual optical properties, whereby each prism acts as an individual optical fiber guiding the ambient light to the inner shell cavity by total internal reflection. This pixelated light channeling enhances both spatial resolution and contrast while reducing angular blurring, an apt combination for acute tracking of a moving object. Our findings may offer insights into the evolutionary aspects of light-sensing and imaging by biological materials and introduce a conceptual framework for the development of bio-inspired multifunctional ceramics and architectured light-tracking materials.

## Main

The microarchitectural organization presented in mineralized shells of mollusks is mainly attributed to the primary function of “protection against external threats” ^1^. Accordingly, the structural complexity arose from coexistence and/or local alteration^2^ of the deployed microarchitectures, such as cross-lamellar, prismatic, foliated, and nacreous, and introduced inter-crystalline and intra-crystalline interfaces are explained as factors serving the mechanical function of the shells. Among these, the nacreous architecture got the greatest attention thanks to its promising fracture toughness^3^. Notably, in the *Pinnidae* family, despite the bivalves’ capability to form the nacreous layer^4^, the posterior side of the shells is solely formed of a prismatic layer^5, 6^. Here, we are reporting an unexpected observation that shows the prismatic layer of the *P. nobilis* shell transmits the environmental light and projects shadows of external objects to the inner part of the shells, integrating a remarkable optical system into the mineralized armor of the animal. We hypothesize that this optical characteristic may be associated with the photobiological behavior of the animal, such as known shell gapping reactions to changes in environmental light intensity^7, 8, 9, 10^ and the shells closure reaction to approaching shadow^7, 10^. Accordingly, we designed and conducted a series of optical experiments to understand the light-shell interactions in different length scales ranging from single prism to full-scale shell and uncovered an optical characteristic possibly related to this function.

## Results

### Translucency

*P. nobilis* – noble pen shells or fan mussels – are shallow water mollusks well-studied for the uncommon geometrical and structural features of their shells^5^, which can grow up to 120 cm^11^. Unlike many other bivalves, *P. nobilis* often stands in a vertical position (Fig. 1a and Supplementary Fig. 1), inserting the tapered anterior ends of the shells in the seabed and anchoring them with the strong byssus threads^10, 12^. While the tapered anterior end of the shell is nacreous - prismatic bi-layered^5^, the posterior side is solely formed by single-crystal-like prismatic calcite (Fig. 1b). The presence of prismatic structure as a dominant fraction of the seashell surface without the second layer is extremely rare, and the presence of single-crystal-like prisms extending from the shells’ outer surface to the inner cavity is only limited *P. nobilis* and *Atrina* (Pterioid bivalves)^13^.

**Fig. 1.**
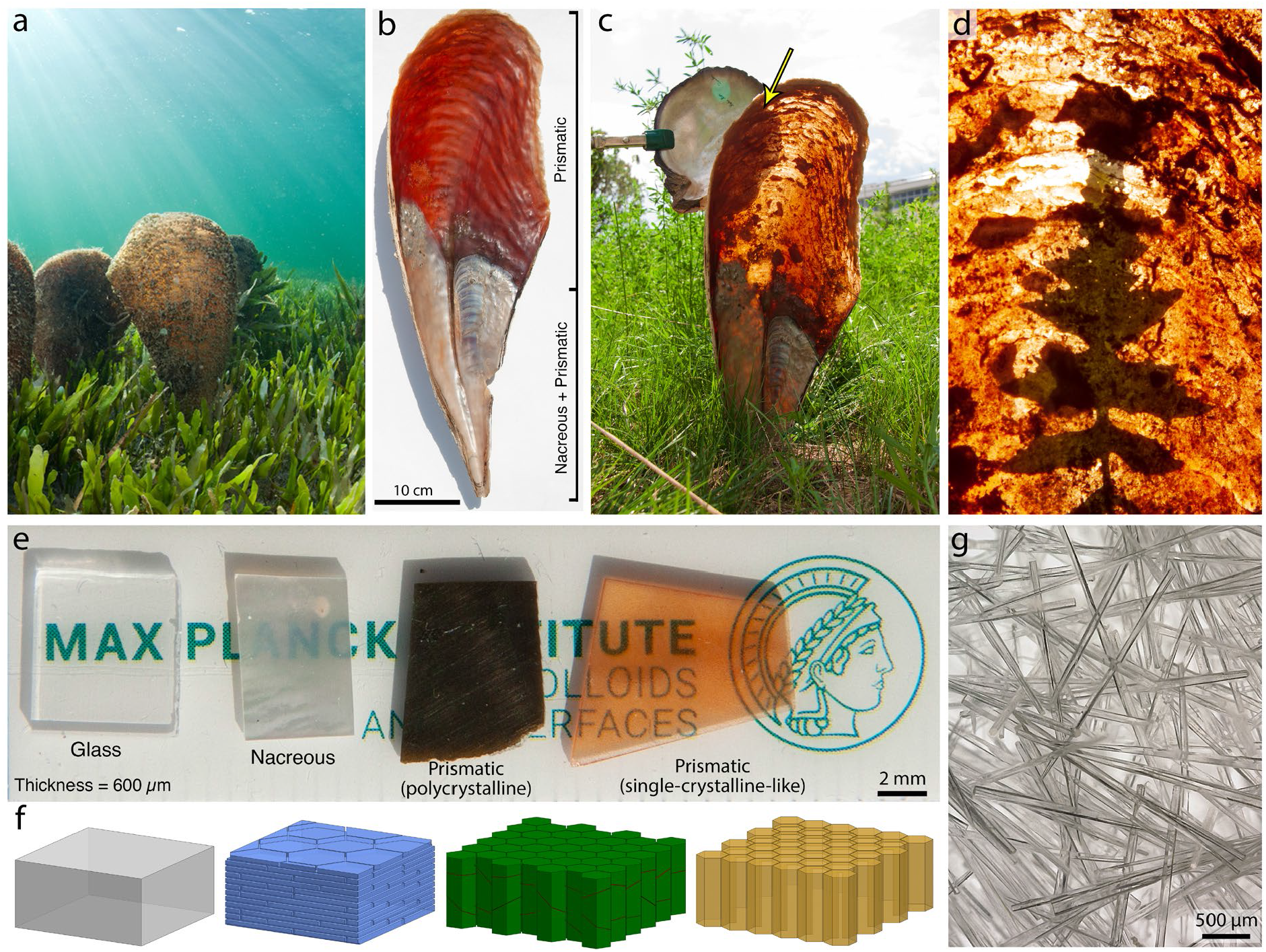
Optical translucency of *P. nobilis* shells. **a**, *P. nobilis* in its habitat (Mar Menor lagoon, Spain) Credit: Javier Marcia. **b**, Photograph of the inner side of a *P. nobilis* shell revealing the absence of the nacreous layer at the posterior part of the shell. **c**, Advances of the light through the entire posterior part of *P. nobilis* shell projecting the shadow of surrounding objects on the inner wall of the shall. In contrast, the *Pinctada* shell could totally block the sunlight (Supplementary Fig. 1 and Supplementary Video. 1). **d**, Photograph highlighting the projected shadow of a leaf with its clear margins on the internal wall of a *P. nobilis* shell. **e**, Photograph of individual calcitic slices (t=600 µm) compared to a glass slide placed on the MPICI institution logo without exposure to any backlight. **f**, Schematic illustration of the corresponding microarchitectural organization of the individual slices presented in panel d. **g**, Single-crystal-like prims of *P. nobilis* shell after treatment by bleaching agent for removal of the organic envelops.

In mature *P. nobilis*, the prisms can reach up to 3 mm (Fig. 1g), extending from the shells’ outer surface to the inner cavity. Surprisingly, despite the shell thickness (prism length), light can advance through the entire prismatic shell and project the shadows of surrounding objects on the inner wall of the shells (Figs. 1c,d, and Supplementary Video. 1). Considering the habitat of *P. nobilis* – shallow water zones of tropical and sub-tropical seas – the animals in their native condition are exposed to environmental light, and their gapping behavior is sensitive to light intensity^8, 9, 10^. Hence, the interaction with environmental light/shadow, even under full moonlight^8, 9^, is considered to contribute to the biologically relevant function of the animals, such as circadian and circalunar rhythms^10^.

Transparency in seashells has been studied in foliated shells of *Placuna placenta*^14^. However, to the best of our knowledge, the transparency in prismatic shells has been noticed^15, 16^ but not been studied. Accordingly, *Pinctada* (*margaritifera)*, a nacro-prismatic model connected to *P. nobilis* from a taxonomical point of view, was selected as a comparative model. However, unlike *P. nobilis*, the outer prismatic layer in the *Pinctada* shell is almost entirely underlaid with a nacreous layer. Notably, by exposing the *Pinctada* shells to ambient light, we noticed that the shell could block it. This was also true when the shell was exposed directly to sunlight (Fig. 1c and Supplementary Fig. 2).

**Fig. 2.**
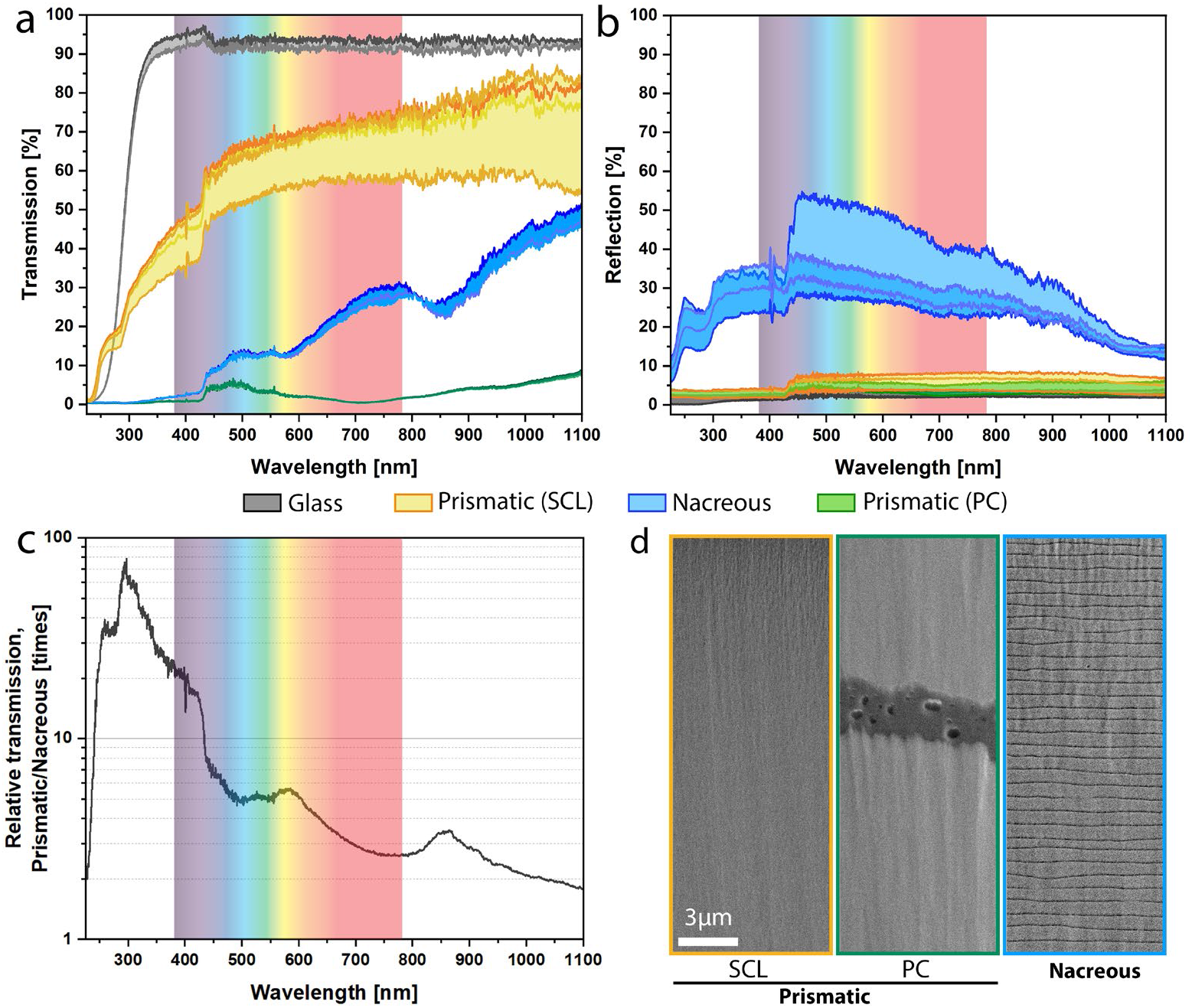
Optical characteristics of individual prismatic and nacreous layers. **a**, Light transmission of *P. nobilis* prismatic shell slice (t= 0.6 mm, depicted in orange) compared with a glass (in grey) and *Pinctada* prismatic shell slice (in green), denoting the high translucency of the *P. nobilis* prismatic shell. **b**, High reflectivity of the nacreous architecture matches the layer’s inability to transmit incident lights. **c**, Relative light transmission of prismatic and nacreous layers presenting 2 to 80 times higher light transmission in the prismatic layer of *P. nobilis*. **d**, Scanning electron micrographs from an ion-sectioned transversal plane of the shells illustrating the absence of the organic interfaces in the single-crystal-like prismatic layer of *P. nobilis* (SCL). In contrast, the polycrystalline prismatic layer (PC) and the nacreous layer of *Pinctada* possess thick and repetitive organic interfaces, respectively.

To understand the light interaction with the individual shell layers and their microarchitectural organizations, we isolated individual shell layers by sectioning and polishing them to form slices with the same thickness (t=600 µm) and compared them with a glass slide as a transparent reference. Figure 1e shows that besides glass, the light transmission through the *P. nobilis* shell slice allowed a clear vision of the text with no distortion or double refraction. We attributed this clarity to the preferred orientation of the calcitic prisms of *P. nobilis* along their *c*-axis^5^ – equal to the ordinary optical axis of calcite – minimizing, if not avoiding, light transmission along the extraordinary axis of calcite and cancels the double refraction^17^. Besides, the single-crystal-like structure of the *P. nobilis* prisms and their extension all along the shell wall result in the absence of interfacial reflection and scattering^17^. In stark contrast, the prismatic slice of the *Pinctada* shells – known for their polycrystalline structure with their *c*-axis perpendicular to the long morphological axis of the prism^18^ – prevented any light transmission. The nacreous slice showed a better light transmission, yet it barely allowed detection of the text.

To quantify light interactions, we measured the relative transmission and reflection of the slides by exposing them to a light source with a broad spectral coverage (250-1100 nm). The collected spectra revealed that the *P. nobilis* slice transmitted the main fraction of the incident visible light (50-70%), while in *Pinctada*, the polycrystalline^19^ prisms and the nacreous layer only transmitted a minor fraction (10-30% and less than 5%, respectively) (Fig. 2a). Notably, the light transmission in the *P. nobilis* shell was not limited to the visible range and included 15-45% transmission in the UV range. In sharp contrast, the nacreous structure acted as a mirror, reflecting 20-50% of the incident light, including UV (Fig. 2b). This mirror-like behavior can be attributed to the architectural organization of the nacreous layer^20^ and the thickness distribution of the platelets (400 - 750 nm), which stays in the ambient light wavelength range (Fig. 2d-right panel and Supplementary Fig. 3). A quantitative comparative analysis was made by plotting the relative light transmission of the prismatic and nacreous samples revealing a 2 to 80 times higher light transmission in the prismatic layer of *P. nobilis* shell (Fig. 2c). Finally, the low (<5%) reflectivity of the polycrystalline prisms along with the low (<5%) transmission indicated that the main fraction of the incident light to this layer was absorbed or internally scattered.

**Fig. 3.**
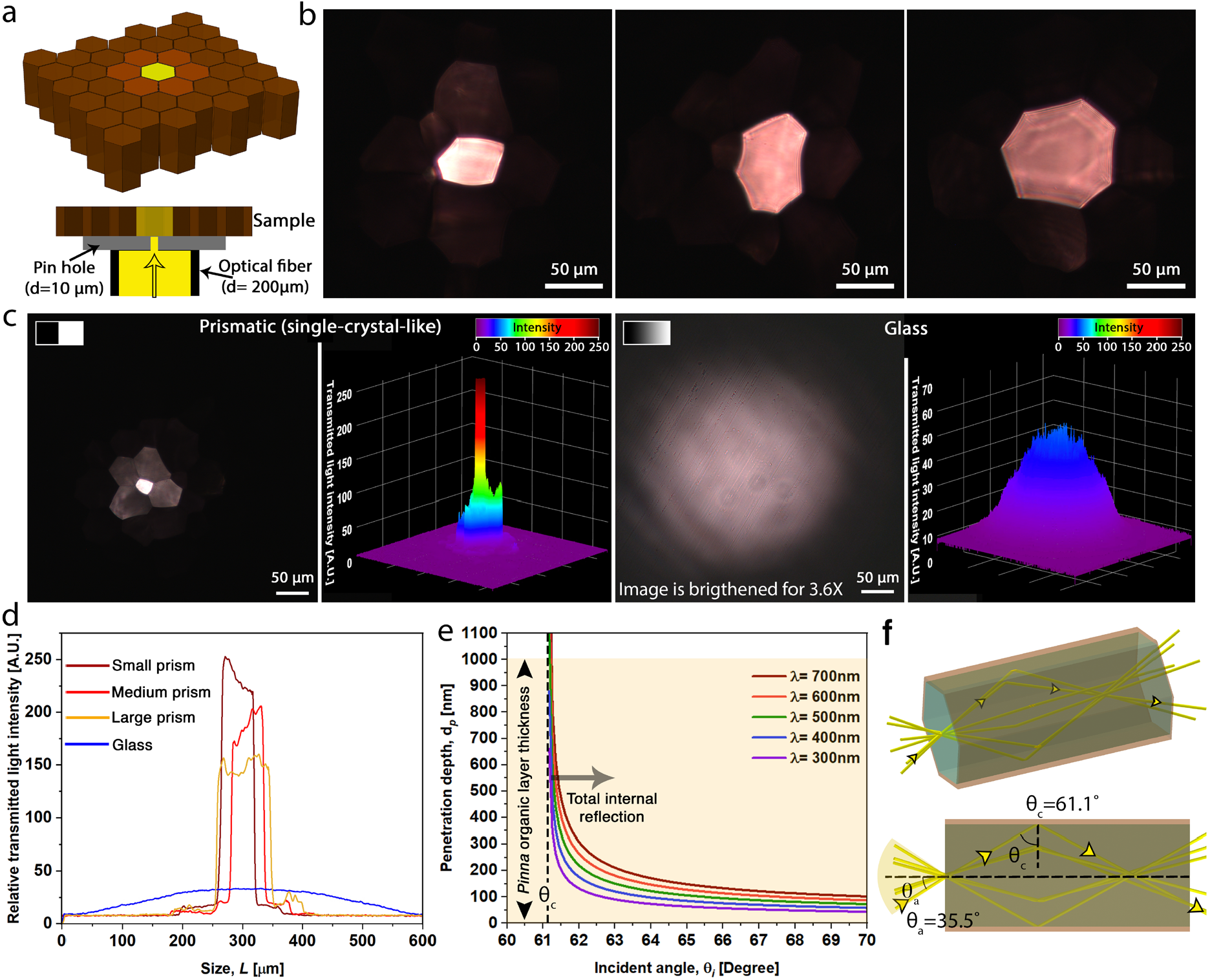
Total internal refraction results in guided transmission of light. **a**, Schematic image of the setup used for intra-prismatic light transmission studies. **b**, Transmitted light through the prisms directly exposed to a 10 µm pinhole revealing that light can be contained within the prisms. **c**, Comparative illustration of the transmitted light profile from the calcitic prism (left panels) and a glass slide (right panels) presenting that while the volume of the transmitted light (measured by volume integration of the light profiles) through the glass is higher, the localized intensity of light in the prism is about 5-fold higher. **d**, Comparative illustration of 2D intensity plots of the transmitted lights in prisms with the different cross-sectional areas and a glass slide. **e**, Calculated penetration depth of evanescent wave for different wavelengths showing that the incident lights to the inner walls of the prisms can barely pass the thick organic envelope (t=1 µm), avoiding the inter-prismatic cross talks. The dashed line represents the critical angle. **f**, Schematic image illustrating the light rays - *P. nobilis* prism interactions and calculated acceptance angle (*θ*_*a*_) and critical angle (*θ*_*c*_) in seawater environment (RI=1.34).

### Containment and channeling of light by total internal reflection

To uncover the light transmission mechanism of the single-crystal-like prisms (d=50-100 µm), we examined *P. nobilis* shell slices (t=1mm) by exposing them to a narrow beam with a spot size of 10 µm (see Methods), ensuring that the light can only enter one prism (Fig. 3a). The observed transmitted light spot with minimal, if no, light leakage to the neighboring prisms revealed that each prism could remarkably contain the incident light and guide it across the slice. By moving the light beam along the slice, we noticed that despite the constant intensity of the incident light, the transmitted light intensity varies inversely with prism size (Fig. 3a, b, d): the transmitted light from a smaller prism showed a higher intensity. Moreover, by comparing the light profiles transmitted from *P. nobilis* shell and glass slices (t=1mm), we found that despite a higher light transmission in glass, as evidenced by volume integration of the light profiles, prisms could avoid light divergence, resulting in a concentrated light spot with ∼5-fold intensity higher than the transmitted light from the glass slide. In addition, we realized that the intensity of the transmitted light has a homogeneous distribution across the prims section, contrasting with Gaussian light distribution that develops in non-architectured and homogenous media such as air and glass. Yet, in some cases, this homogeneity in transmitted light intensity experienced an interference pattern (Fig. 3b and Supplementary Fig. 4a), denoting the formation of phase shifts known to be induced by internal refraction inside the prisms^17^. These observations suggested that the transmission of the light inside the single-crystal-like prisms occurs through a total internal reflection phenomenon.

**Fig. 4.**
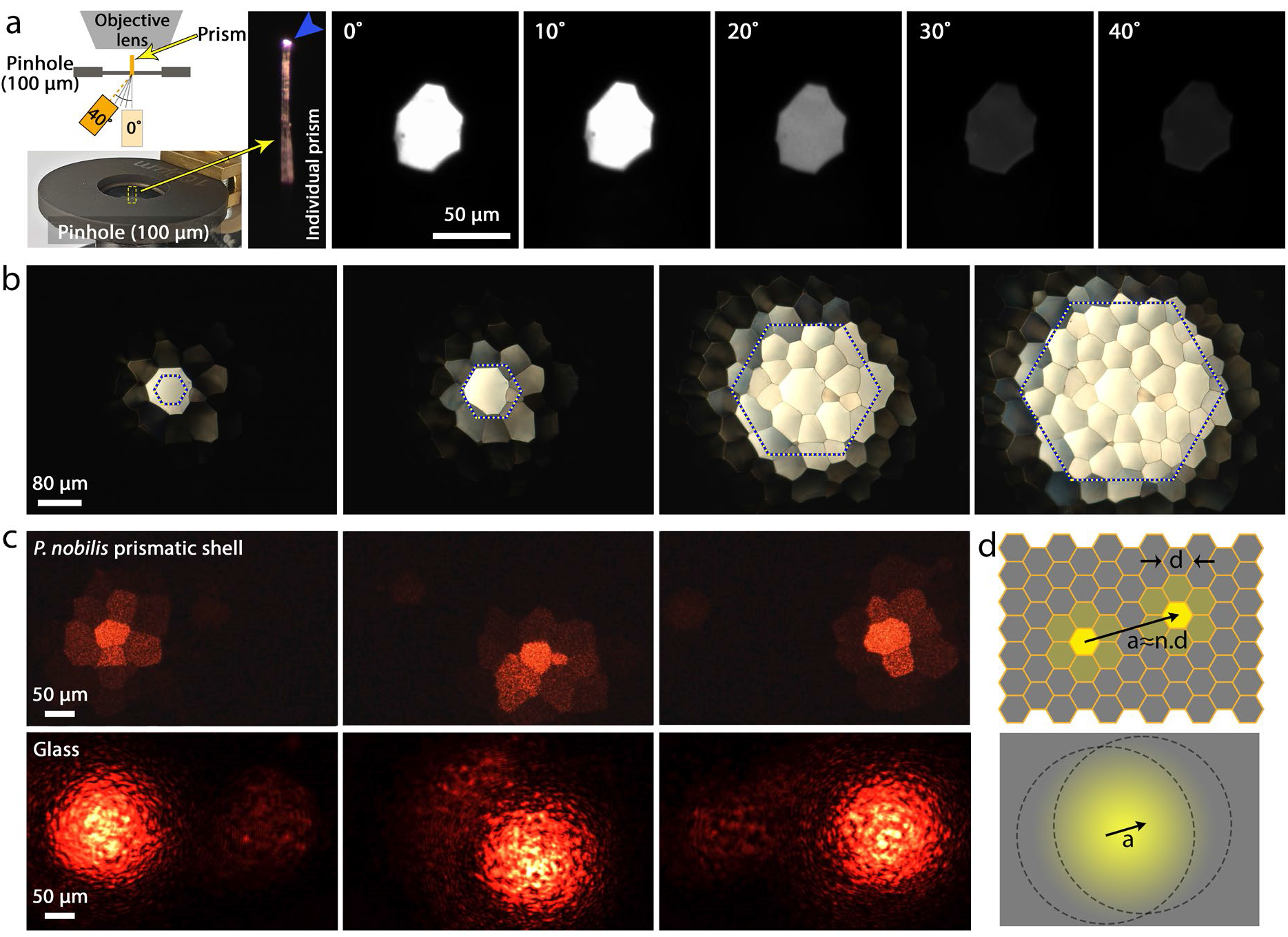
Pixelated containment of light within *P. nobilis’s* prisms. **a**, Guided light through an individual *P. nobilis* prism (l≈1.8 mm) in different angular exposure resembling an optical fiber. **b**, Reflected light spots on the *P. nobilis* slice revealing the mismatches between the peripheral edge of the illuminated prisms and the hexagonal geometry of the light spot formed by the microscope aperture denoted by the dashed hexagons. The microscale mismatches induced by channeling the light within the calcitic prisms provide an accurate fit between the projected and transmitted light/shadow in the macroscale (Figs. 1d,e and Supplementary Fig. 2, middle row). **c**, Transmitted light spots formed by sweeping a light beam along a native prismatic *P. nobilis* shell (upper row) and a glass slide (bottom row), revealing the spatial resolution and contrast in the prismatic *P. nobilis* shell compared to the glass sample (see Supplementary Video 3). This characteristic, formed by the total internal reflection of the light in the calcitic prisms, introduces the prismatic *P. nobilis shell* as an optical light guide array for high-precision light/shadow tracking of a moving object. **d**, Schematic illustration presenting how pixelated light can foster spatial resolution for tracking a moving object.

To better understand the principle of the light interactions within individual prisms and quantify the optical parameters of the *P. nobilis* shell, we used optical diffraction tomography (ODT), a holographic microscopy technique, to quantitatively measure the three-dimensional refractive index (RI) distribution of prism and organic shield (See Methods) individually^21, 22, 23^. As expected, due to the preferred orientation of the calcitic prisms along their *c*-axis (ordinary optical axis), a high RI of 1.61 (n=10, individual prisms) was measured for the calcitic prisms (Supplementary Fig. 5). This value was slightly lower than the reported value for the RI of geological calcite along their optical axis (RI_or_=1.66). This difference can be attributed to the presence of organic residues in the biological calcite^24^. The measured RI value of the organic shield (RI=1.41, n=10, individual pieces of organic shields) and the proximity to the refractive index of calcite along the extraordinary axis (RI_ex_=1.49) revealed how a preferred alignment along the ordinary optical axis (*c*-axis) of the prisms can be beneficial for decreasing the total internal reflection angle. We found that by alignment of the calcitic prisms along their *c*-axis, the critical angle (*θ*^*c*^) of the prisms in seawater (RI= 1.34^25^) decreased by 14% (71.1° in the extraordinary optical axis of calcite vs. 61.1° in *P. nobilis*), resulting in a 68% increment (21.1° in the extraordinary optical axis of calcite vs. 35.5° in *P. nobilis*) in the acceptance angle (*θ*^*a*^)(Fig. 3f). Remarkably, the enhancement in acceptance angle and numerical aperture (NA) resulted in a 2.75-fold increment in light-gathering power (See Methods). In other words, the alignment of the prisms along their *c*-axis can increase the oblique angles to collect a larger quantity of ambient light, enhancing the light sensitivity of the prismatic shells, critical criteria for light interacting models in aquatic environments^26, 27, 28, 29^. It has been reported that the animal can react to sudden darkness even at night under full moon light^8, 9^ implying the high responsiveness of the animal even under low light conditions. This behavior can explain the need for high light-gathering power.

**Fig. 5.**
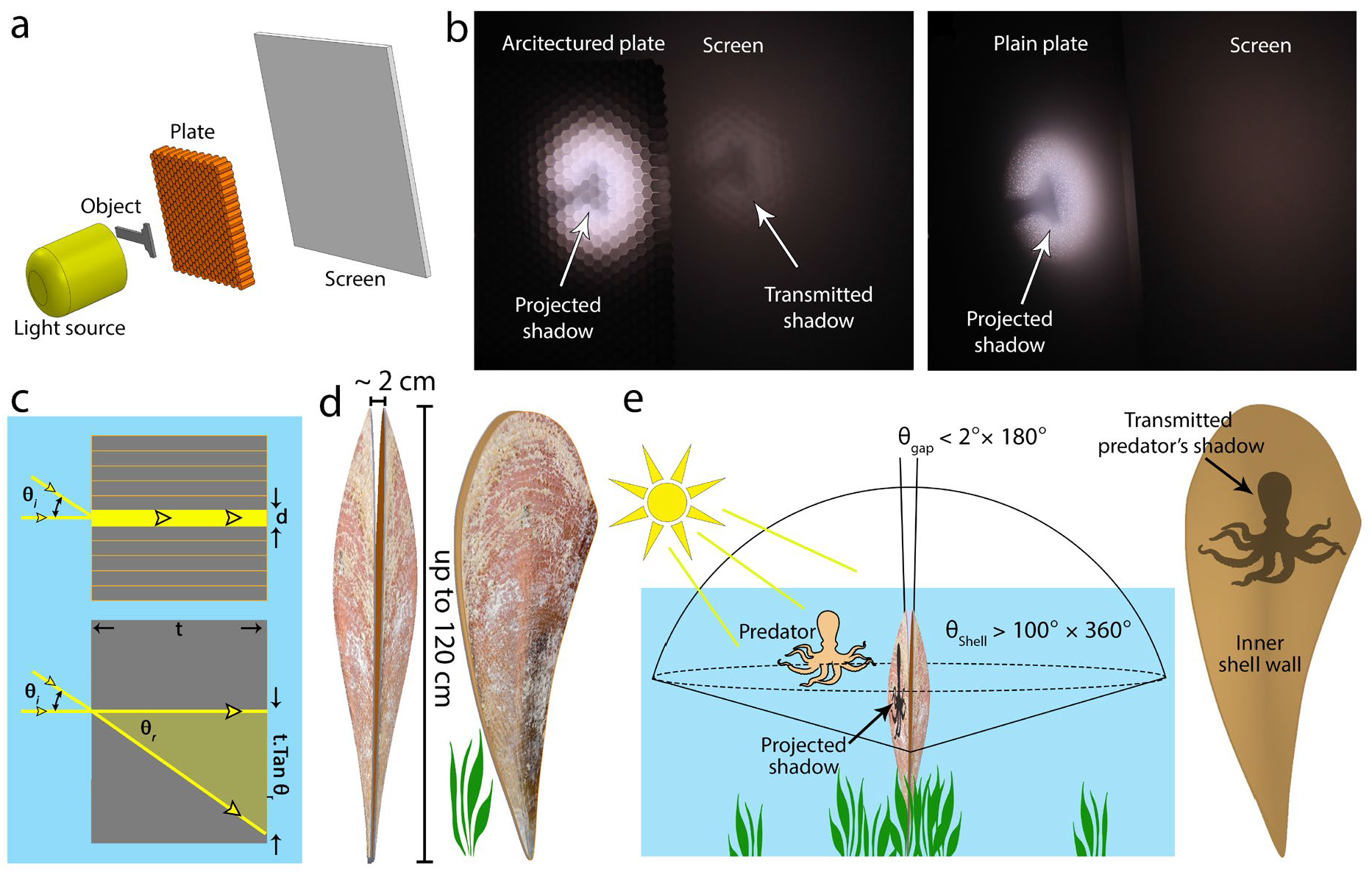
Shadow transmission through prismatic architecture. **a**, Schematic illustration of shadow projection setup used for studying the formation of reflected and transmitted shadows. **b**, Projected and transmitted lights on the architectured and plain acrylic plates. The prismatic light guides of the architectured plate reduce the refraction drifts of the projected light spot on the screen resulting in the formation of a transmitted shadow on the screen. In contrast, the refraction in the plain panel prevents the formation of the transmitted shadow (Supplementary Fig. 7). **c**, Schematic illustration presenting how the prismatic light guides promote a reduction in sensitivity to the incident angle, retain the incident light/projected shadow position, and result in the transmission of the shadow with high contrast and spatial resolution. **d**,**e**, Considering the narrow shell gap (minute light window), transmitting light/shadow through the prismatic layer can confer wide-angle light access on the animal independent of the sun/moon position.

In addition, the calculated penetration depth of evanescent waves^17^ for different wavelengths of the incident light presented the importance of a thick (1-2 µm) organic envelope in preventing light leakage to the neighbor prisms. Nevertheless, in some cases, we could detect light leakages to the neighbor prisms (Fig. 3c-left panel and Supplementary Fig. 4b). We suggest that the size and taper angle of the shrinking or growing prisms^30^ may violate the critical incident angle and contribute to the leakage (if any) of the contained light to the neighbor prisms. Later, we designed an experiment to evaluate the performance of an individual – isolated – calcitic prism (with a prism length l=1.8 mm) by exposing it to a light beam (with optical fiber diameter d=200 µm) at different incident angles (Fig 4a, see Methods). In good agreement with our previous observations, the transmitted light at the free end of the prism showed a bright glow (Fig 4a, blue arrow), closely resembling the function of an optical fiber/ light guide. Our observations showed that despite the changes in the intensity of the transmitted light, by changing the incident angle, the light can be fairly transmitted through the individual prism.

### Enhancement in spatial resolution and contrast

Because of the displayed optical characteristics and geometrical features of the individual calcitic prisms of *P. nobilis*, we compared them with the apposition compound eye’s crystalline cones^31, 32, 33^, in which cones act as optically isolated light guides guiding the light by total internal reflection^32, 34, 35^. Hence, the physical characteristics of the cones bundle define the resolution of the spatial information which can be transferred to the compound eye. Accordingly, to examine the optical performance of the calcitic prisms and evaluate the interrelationships between the microarchitectural organization of the prisms and induced spatial resolution, we designed two experiments in reflection and transmission modes.

By probing *P. nobilis* shells with a reflected light microscope and changing the light spot size with the microscope aperture, we noticed that the shaped light spot on the shell did not match the hexagonal geometry of the aperture (Fig. 4b, dotted hexagons). Instead, the light spot was defined by the basal size and arrangement of the illuminated prisms (Fig. 4b), such that even by exposure of a small fraction of one prism, the entire prism was uniformly illuminated and outlined a new perimetric border. This denoted that each calcitic prism acts as a binary pixel that, upon the exposure or closure of the light, can be switched ON or OFF. Hence, by changing the light spot size, resembling a moving object, the perimetric prisms present a clear and acute track of light development. It means that the *P. nobilis* prismatic shell acts as an apposition light guide array and can convey the spatial information of the incident light with resolution down to tens of micrometers, which is the basal size of the prisms (Fig. 4b and Supplementary Video 2).

We then investigated the role of prismatic architecture and induced spatial resolution in tracking a moving object using transmitted light and whether this feature can be preserved across a native *P. nobilis* shell. Accordingly, we exposed a native *P. nobilis* shell (t ≈ 2mm) to a light beam (laser diode, λ= 650 nm) while moving the beam along the sample surface (Fig. 4c, upper row). A glass slide (t= 1mm), representing a translucent plain sample, was selected as a comparative reference (Fig. 4c, lower row).

The pattern of the illuminated prisms indicated that by sweeping the light source along the shell surface, *P. nobilis* prisms could form a binary (ON-OFF) illumination (Fig. 4c, upper row, and Supplementary Video 3), which retains and transfers the spatial resolution and contrast of the external light/shadow spot to the inner surface of the prismatic shell. The exhibited resolution and contrast can potentially provide the required information for acute positioning and estimating the speed of a moving object’s shadow (Supplementary Note 1). These outstanding features cannot be attained by homogenous media independent of material type (Fig. 4d).

### Reduction in sensitivity to angular exposure

We extended our investigations on the role of the prismatic array and induced pixelated light channeling on the formation of reflected and transmitted lights/shadows by designing and conducting an experiment on a replicated model. Accordingly, a frosted acrylic plate (t= 3.2 mm) was laser cut (see Methods) to form an array of hexagonal prisms (Fig. 5). By exposing the architectured plate to a white light source and comparing it with a plain acrylic plate, we realized that, firstly, the formation of a hexagonal light pattern resulted in a clearer light contrast on the architectured plate compared with the plain plate (Fig. 5a). Secondly, thanks to a guided transmission of light through the prisms, the light intensity and shadow contrast were retained and transmitted as evidenced by the preserved hexagonal pattern on the screen. In contrast, the light divergence amplified by the refraction of the plain plate did not allow a clear formation of the transmitted light spot, resulting in the blurring of the object’s shadow.

Moreover, we hypothesized that the architectural organization of the prisms could lower the sensitivity to the incident angle of the light creating the image on the shell surface. Accordingly, we tested our hypothesis by exposing the plates to incident light at different angles (Supplementary Fig. 7a,b). Remarkably, we found that the guided light transmission induced by the prismatic light guides can reduce, if not cancel, the drifts in the position of the projected light spot, resulting in the reduction of the angular sensitivity for better spatial positioning of the light/shadow. In contrast, the transmitted light spot through a homogenous sample presented an obvious angular-dependent deviation from the incident point regulated by the RI and thickness of the plate (Fig. 5c and Supplementary Fig. 7b,c).

## Discussion

In summary, our observations provided empirical evidence of sophisticated light guide arrays integrated into the mechanical armors – shells – of *P. nobilis*, combining unlike and conflicting functions and devising a multifunctional structure^29, 36^. We showed how the individual calcitic prisms act as optical fibers retaining and transferring the environmental light through total internal reflection to the inner shell cavity. We revealed that the bundled prisms – prismatic shell – form a light guide array converting the external light/shadow profile from a light intensity gradient to a pixelated intensity map and projecting it on the inner shell surface. This conversion enhances the spatial resolution (defined by the prism diameter) and improves the contrast (achieved by sharp light intensity differences between neighboring prisms), yielding an optical feature suitable for the tracking of moving shadows. Notably, a similar pixel-array mechanism is being deployed in technological applications dedicated to image transferring^37, 38, 39, 40^ or pre-processing spatial light information for advanced motion-detection, edge-detection, and orientation-detection devices^41, 42, 43^. This pre-processing implemented in the hardware through the pixel-arrays reduces the complexity of the subsequent computational image processing^43^.

It is not clear, whether these exceptional optical characteristics of the *P. nobilis* prismatic shell correlate in any way with the animal’s biological behavior. However, the reaction of *P. nobilis* to changes in environmental light intensity is documented^8, 9, 10^, and there are occasional reports that the animal closes its valves with a shadow approaching, which led to the suggestion that *P. nobilis’* mantle possesses a light-sensing mechanism^7^. Indeed, the closure of the valves during the approach and the presence of predators can be considered an essential precondition for effective protection. Considering the upright standing position of the animal (Supplementary Fig. 1) and their limited gaping aperture (∼2 cm)^8, 9^, the shell gap only opens a narrow notch, which does not allow much access to surrounding light or shadows. In contrast, the translucency of the shell through the light guide arrays (prismatic bivalves) opens up a window with wide-angle visibility and significantly expands the animal’s access to light and shadow of surrounding objects (Fig. 5d,e). In addition to mechanical protection, light-sensing is known to play a role in the circadian clock of the animal^10^.

However, it remains to be proven that the revealed light guide array of the *P. nobilis* shell is part of a light-sensing or light-harvesting system to provide shadow tracking or imaging^32, 44^. We are also aware that invasive marine organisms, such as biofouling in the Mediterranean Sea, can influence these optical characteristics. However, we reason that considering the size of the immense numbers of the prisms, partial coverage of the shell by external objects does not block the access of the shell to the environmental light.

We propose that our findings may offer insights into the evolutionary aspects of light-sensing and imaging by biological materials and introduce a conceptual framework for the development of multifunctional transparent ceramics^45, 46^ and architectured light-tracking materials.

## Methods

### Sample collection

Two groups of *Pinna nobilis* and *Pinctada margaritifera* shells were used for the studies. The full-size shells were borrowed from the Naturkunde Museum of Berlin for photography, as depicted in Fig. 1 a-c). A series of wild smaller *P. nobilis* from the Mediterranean Sea and *Pinctada* from French Polynesia shells were used for material and optical studies.

### Sample preparation

Preparation of slices: To prepare the shell slides, *P. nobilis* shell pieces were directly polished using abrasive papers and colloidal particles down to 1 µm. For *Pinctada* samples, the nacreous and prismatic layers were first separated along the interfacial plane using a diamond wire saw (d=150 µm). Then, each layer was individually polished down to 1 µm. Preparation of Individual prisms: Individual *P. nobilis* prisms were prepared by immersing small (∼5 mm x mm) shell pieces in a bleach solution (Solidum Hypochlorite) for 24 hrs at room temperature. Preparation of Optical diffraction tomography sample: Individual *P. nobilis* prisms from the prismatic layer of the pinna nobilis shell were fixed along the *c*-axis of calcite with DI water under -20°C then cut into slices with a thickness of around 30 µm using a Leica Microtome (Leica CM3050 S, Leica Microsystems Nussloch GmbH, Germany). To separate the organic shields, small *P. nobilis* pieces were immersed for 48 hrs in a commercial acidic bleach solution (PH=1.2, Ja Detergent Tab) at room temperature to bleach the calcitic prisms (Supplementary Fig. 8). The bleached sample was rinsed and later sonicated in water for 5 minutes to separate the organic shields. Small droplets containing the organic shields were placed on the glass coverslips and were dried under a fume hood. Preparation of polyacrylic samples: The architectured replicated polyacrylic panels (t= 2 mm) were designed and later outsourced for laser cutting. The pristine polyacrylic plates were used as a comparative reference.

### Optical microscopy

The optical images used in Fig. 1b-e, Fig. 5b,d, and Supplementary Fig. 2 were captured using a DSLR camera (EOS-500D, Canon) equipped with an EFS 18-200 lens (Canon). The optical micrograph of individual prisms (Fig. 1g) was captured using a stereomicroscope (Leica-DVM6). Optical micrographs were captured using a microscope (Zeiss, AxioLab 5, Germany). For the studies in Fig. 3b,c, an external light source (DH-2000-BAL, Ocean Insight), an optical fiber with a diameter of 200 µm, and a pinhole (d=10 µm, Thorlabs) were used to provide the transmission light. The slice samples were placed directly on the top of the pinhole, and the transmitted light from the sample was collected using a 20X objective lens (Zeiss).

### Light transmission and reflection studies

The light transmission and reflection studies were performed using a customized setup including a light source (DH-2000-BAL, Ocean Insight), a spectrometer (OCEAN-HDX-XR, Ocean Insight), and a set of optical cables (One Y-type fiber for reflection studies and two normal fibers for transmission studies). The collected spectra were extracted using OCEANVIEW (Ocean Insight) and then plotted by OriginPro 2020.

### Ion-sectioning

Ion-sectioning of the shell layer slices was done using a cross-section polisher (IB-19520CCP, JEOL) for 5 hrs of course cut (accelerating voltage 5.5kV) followed by 1.5 hr fine polishing (accelerating voltage 3kV). To decrease/prevent the thermal damage to the organic residues, the sectioning was done in cryo-mode (T= -120°C), and a cutting cycle of 40 s ON – 20 s OFF was set for the cutting.

### Electron microscopy

Ion-sectioned samples were probed using a Field Emission Electron Microscope (FESEM, JEOL 7200) at an accelerating voltage of 5 kV. The samples were coated using carbon for ∼10 nm prior to the study.

### Light interaction studies on individual prisms

The studies on the individual P. nobilis prims were done under a stereomicroscope (Olympus, SZX7) equipped with an sCMOS camera (Andor, Zyla). The individual prism was inserted inside a pinhole (d= 100µm), and the griped end was positioned in the rotation center of a swivel mount (Thorlabs). A spotted light using an optical fiber (d= 200 µm) was exposed to the gripped end of the prism, and the transmitted light was captured from the free end of the prism at the incident angle of 0°, 10°, 20°, 30°, and 40°. The images were captured using the camera software (Andor Solis).

### Shadow projection

The transmission of projected shadow (Fig. 5b) was investigated using a white light source (CoolLED, pE-4000) and a viewing screen (Thorlabs, EDU-VS1).

### Optical diffraction tomography (ODT)

The three-dimensional (3D) refractive index (RI) of isolated calcitic prisms (thickness 20–30 µm) and unfolded organic shield layers (thickness 1–3 µm) was measured using a custom-built optical diffraction tomography (ODT) microscope employing Mach-Zehnder interferometry similar to the one described in Ref (^21^). The Mach-Zehnder interferometry measured spatially modulated holograms from 150 different angles, from which the complex optical fields were retrieved. The 3D RI tomograms were reconstructed by mapping two-dimensional Fourier spectra of retrieved complex optical fields onto the surface of the Ewald sphere in the 3D Fourier space according to the Fourier diffraction theorem. Detailed principles for tomogram reconstruction can be found in Refs (^22, 23, 47^). To image the prism and the organic layers, thin sections of each were sandwiched between two glass coverslips (VWR, Cat. No 631-0165). In order to reduce the RI mismatch between the samples and surrounding media, the prism layer was embedded in oil of RI_oil_=1.6100 (Series A Cargille Laboratories, Cat #: 1809), and the organic layer was embedded in oil of RI_oil_=1.3950 (Series AAA Cargille Laboratories, Cat #: 1803). The image acquisition, field retrieval, and RI tomogram reconstruction were performed using custom-written MATLAB scripts (R2020a). The mean RI value of each sample was measured with FIJI^48^ and the statistical analysis was performed using GraphPad Prism.

### Data analysis

Data analysis and plotting were done using OriginPro software. Values are reported in mean+/- SD format.

## Data availability

Data are available from the authors upon request.

## Acknowledgments

We thank J.P. Cuif for his generous help in providing the *Pinna nobilis* and *Pinctada* (*margaritifera*) shells samples and Jochen Guck for his support for RI mapping measurements. We thank Oliver Späker for his help with RI measurements. We acknowledge C. Zorn (Museum für Naturkunde, Berlin) for supporting us by providing the *Pinna nobilis* and *Pinctada* (*margaritifera*) full-scale shells. We thank James C. Weaver for introducing the relative technological applications based on fiber optic bundles.

## Author contributions

S.A. and P.F. designed and developed the study. S.A., P.F., and M.C. designed the experiments. S.A. and T.Z. conducted the optical measurements. Y.D. provided the shell samples and insights regarding the studied models’ biological characteristics. A.B., K.K., and S.R. conducted and analyzed the RI mapping studies. S.A. wrote the manuscript with input from all the authors.

## Funding

The project is funded by Max Planck Society. S.R. and A.B. acknowledge funding by the DFG (RE 3925/1-1) and Joachim Herz Stiftung, respectively.

## Competing interests

The authors declare no competing interests.

## Additional information

Supplementary information is available for this paper.

## Notes

https://www.dropbox.com/sh/nn1whwxg8lmev69/AAAIMTirjd0HN7cYa1ZuXVCIa?dl=0

### Competing Interest Statement

The authors have declared no competing interest.

