## Supplementary information for "Calcitic prisms of the giant seashell *Pinna nobilis* form light guide arrays"

### Supplementary Figures 1-8

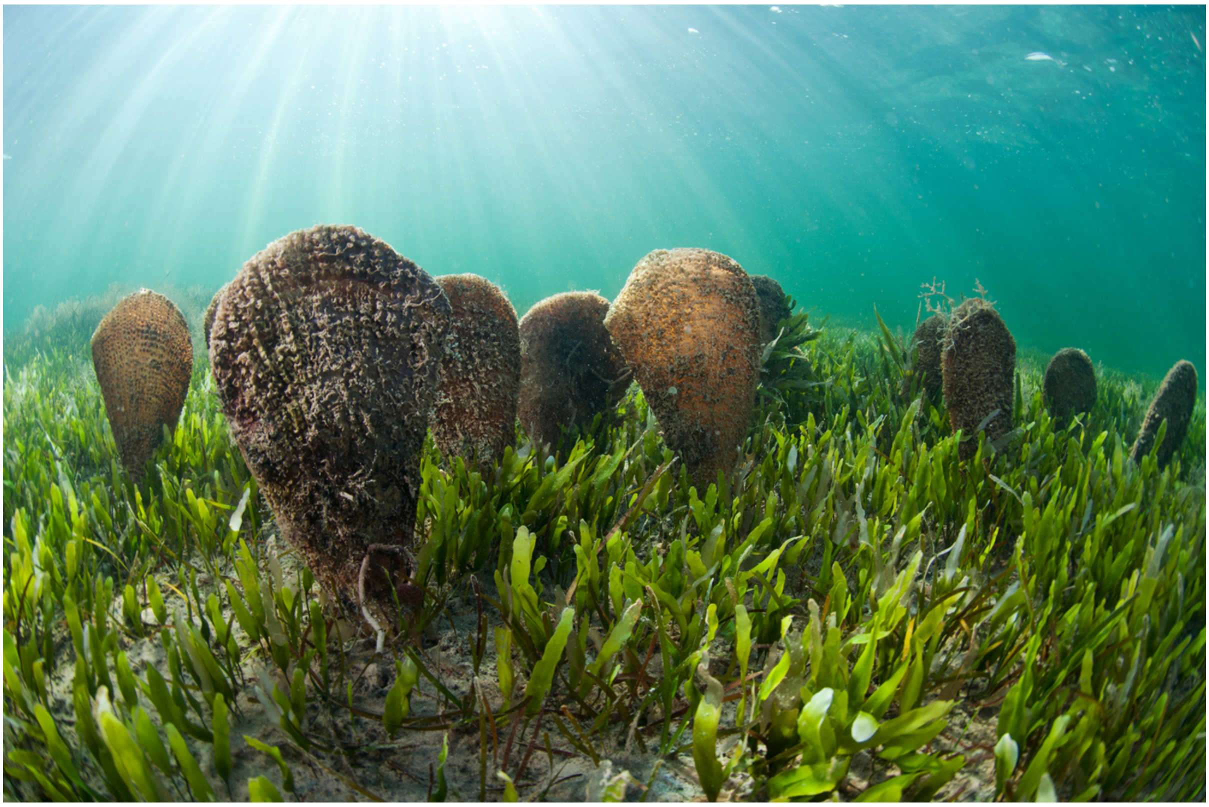

**Supplementary Fig. 1| *P. nobilis* in its habitat (Mar Menor, Spain).** Optical micrograph illustrating the vertical positioning of *P. nobilis*. The animals insert the anterior end of the seashells in sandy and muddy seabeds and anchor them using their byssus threads (Credit: Javier Murcia).

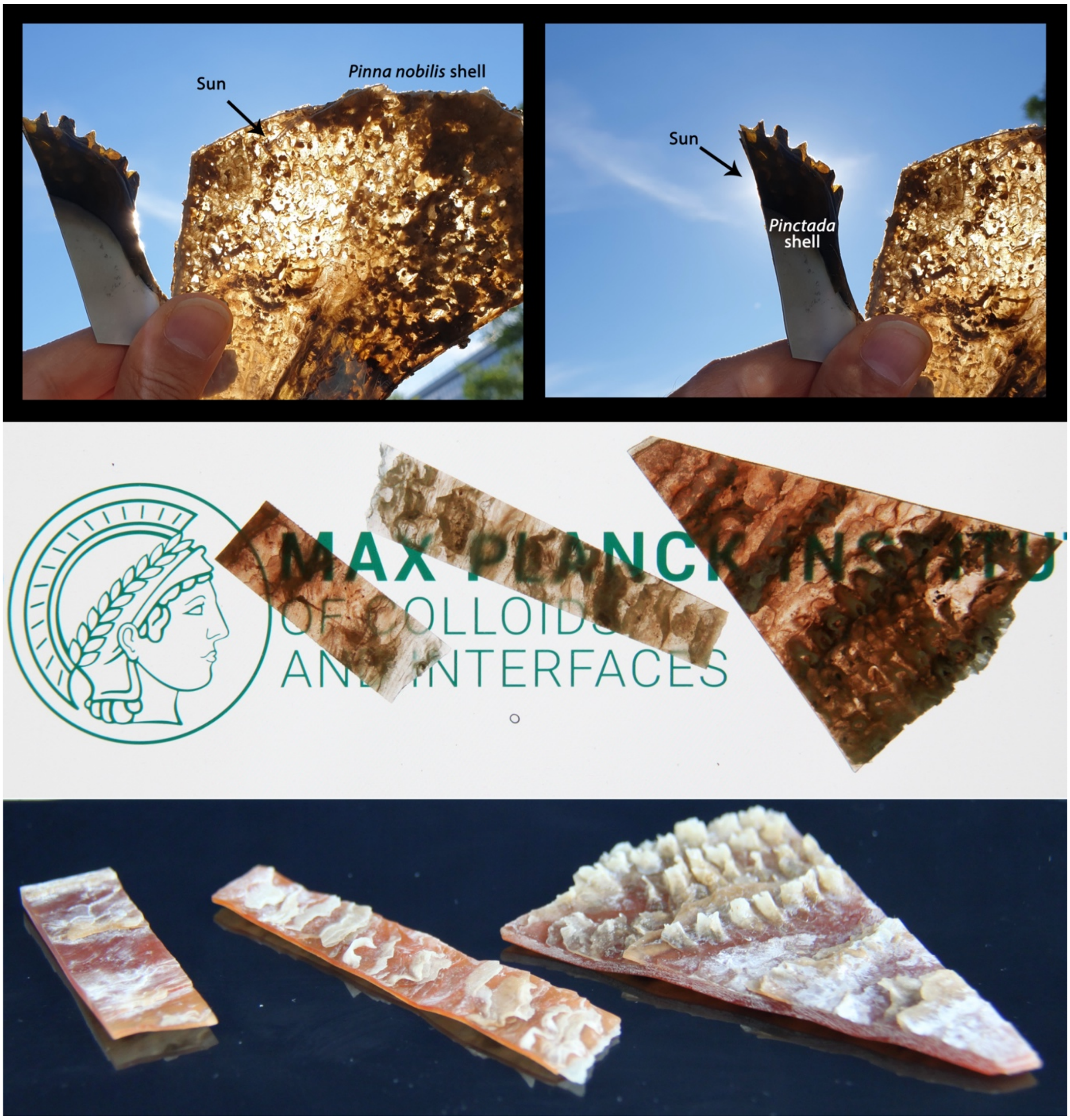


**Supplementary Fig. 2| Optical translucency and obstruction in *P. nobilis* and *Pinctada* shells.** Optical micrographs showing the light transmission of the *P. nobilis* shell compared with light obstruction of the *Pinctada* (*margaritifera*) shell, which can even totally block direct sunlight. Image of native (not polished or bleached) pieces of *P. nobilis* shell on a cell phone screen (with backlight) presenting the translucency of the shells. Bottom row showing the *P. nobilis* shells with no backlight.

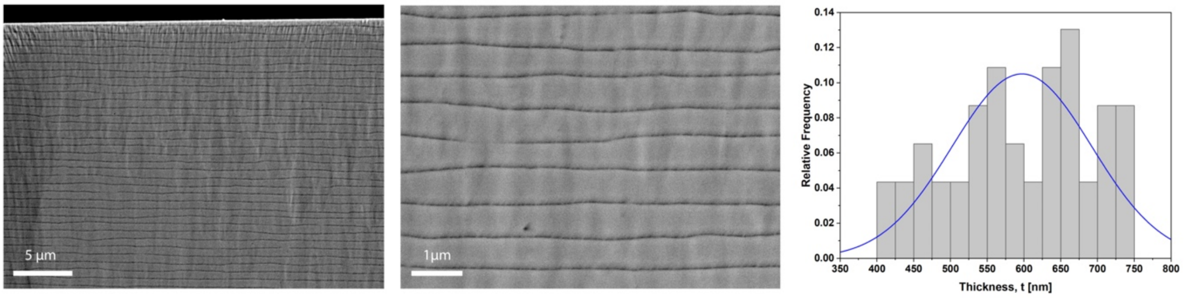


**Supplementary Fig. 3| Distribution of the layer thickness in the nacreous layer of *Pinctada* shell**. Electron micrographs obtained from the cross-sectional plane of a *Pinctada* nacreous layer were used to calculate the distribution plot of the layer thickness. Light reflectivity (Fig. 2b) of the nacreous shell is attributed to the ultra-architecture and size and spacing of the nacreous platelets compared to the visible light wavelengths (~400-700 nm).

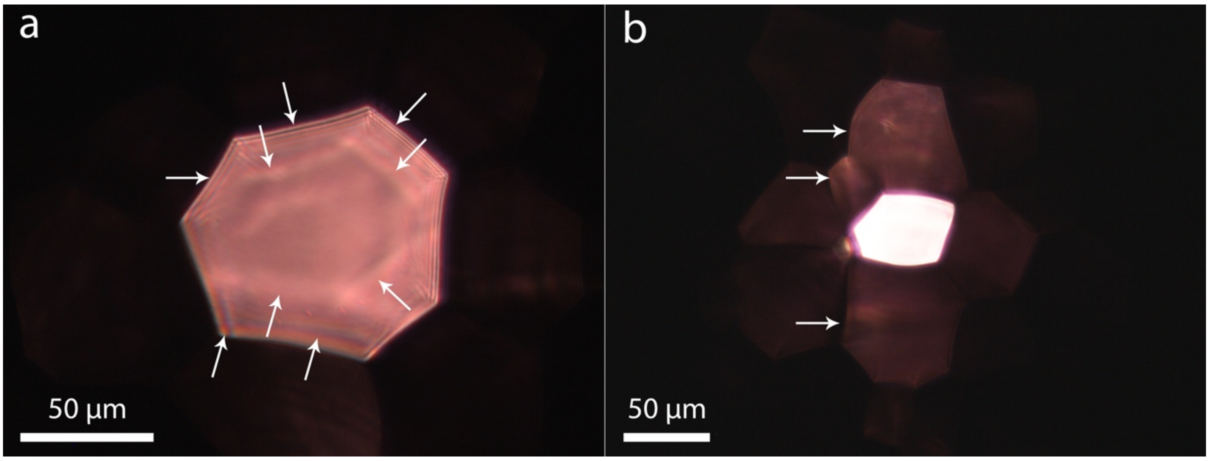


**Supplementary Fig. 4| Interference pattern and light leakage in the *P. nobilis* prisms. a,** Optical micrograph of transmitted light through a *P. nobilis* prism revealing the formed interference pattern, denoted by the white arrow. This pattern is formed by the phase shifts occurring by total internal reflection of the light within the *P. nobilis* prisms. **b,** Optical micrograph of minor light leakages to the neighbor prisms showing the possibilities of the cross-talks between the prisms. The leakage, which only happened in a small fraction of the prisms, might be attributed to the size and taper angle of the shrinking or growing prisms.

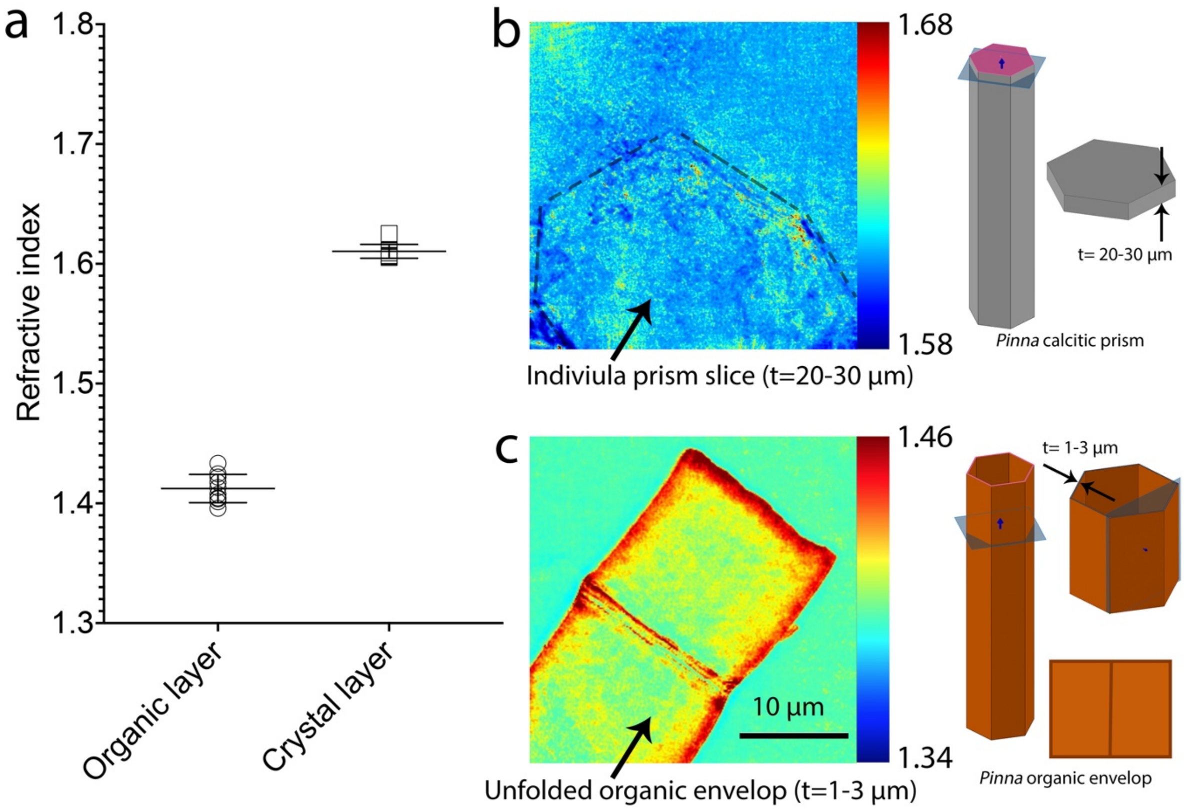


**Supplementary Fig. 5| Refractive Indices (RI) of calcitic prisms and organic envelopes. a,** RI values of calcitic prisms and unfolded organic envelopes calculated from Info on statistics. **b-c,** Representative central slices obtained via ODT showing the RI distribution in (b) an isolated calcitic prism and (c) a layer of organic shields (see Methods).

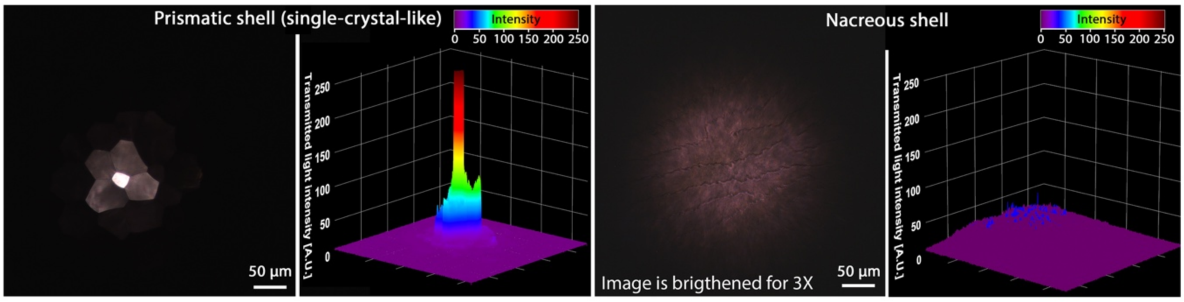


**Supplementary Fig. 6|Containment of light in a calcitic prism of *P.nobilis* shell.** Comparative illustration of the transmitted light profile from the calcitic prism (left panels) and a nacreous layer (right panels) presenting higher light transmission and the localized intensity of light in the prismatic sample.

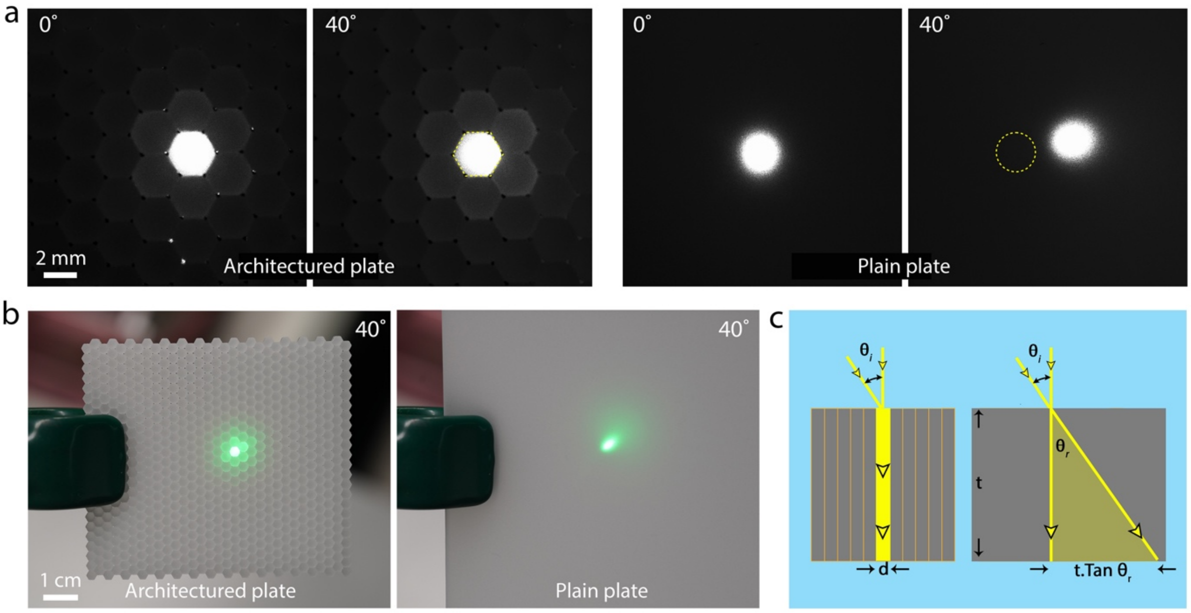


**Supplementary Fig. 7|Containment of light in architectured and homogeneous models at angled incident light. a,** Transmitted light spots from the architectured (left) and plane (right) panels exposed to a white light beam at 0° and 40° incident angles. The architecture plate can retain the position of the incident light across the panel. **b,** Reflected light spots on the architectured (left) and plain (right) plates exposed to a laser beam (λ=532 nm) at 40° incident angle. **c,** Schematic illustration presenting how the prismatic light guides promote a reduction in sensitivity to the incident angle, retaining the incident light position resulting in enhancement of the contrast and spatial resolution under angular exposure.

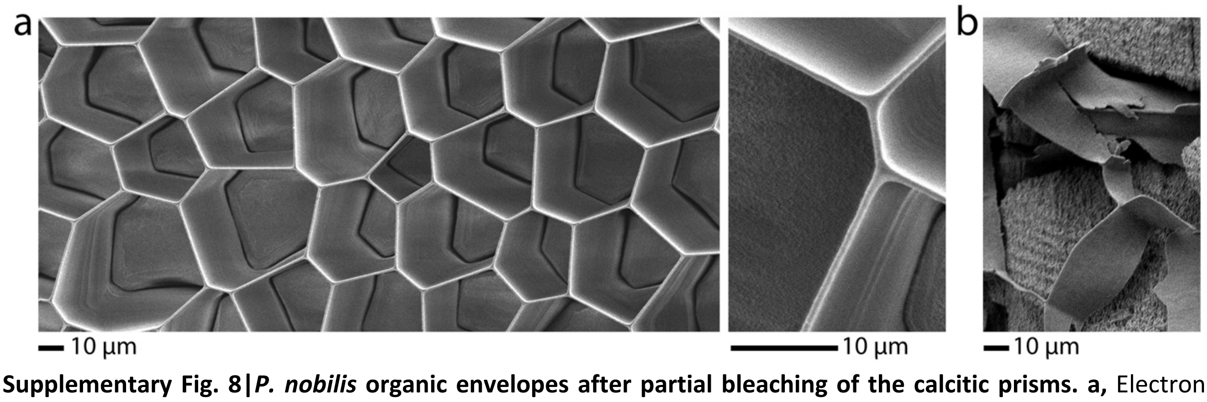


**Supplementary Fig. 8|*P. nobilis* organic envelopes after partial bleaching of the calcitic prisms. a,** Electron micrographs of a *P. nobilis* shell revealing the organization of organic envelops. **b,** Organic envelopes were separated from the shell by sonication of the bleached shell in water.

### Supplementary Videos 1-3 (captions)

**Supplementary Video. 1| Transmitted shadow of the environmental objects (surrounding plants) on the inner wall of the *P. nobilis* shell.**

**Supplementary Video. 2| Development of the reflected light spot on the *P. nobilis* prismatic shell.** Despite the hexagonal geometry of the projected light, shaped by the microscope aperture, the geometry of the formed light spot is defined by the basal size and packing of the calcitic prisms. This characteristic allows the *P. nobilis* shell to provide acute spatial information regarding the advancing light/shadow spot.

**Supplementary Video. 3| Transmitted light spot on the inner wall of a native *P. nobilis* shell (t=2mm), frosted glass (t=1mm), and glass slide (t=1mm).** The light beam (laser diode, $\lambda$=650 nm) was moved along the *P. nobilis* shell (t=1-38s), frosted glass (t=39-57s), and glass slide (t=58s-85s).

**Supplementary Note 1**

**Prismatic light guide array: a measuring tool for quantitative tracking of moving objects**

By exposing the *P. nobilis* prismatic shell – light guide array – to the shadow of an external object, the prisms behave as binary (ON-OFF) switches. This characteristic can be used as a measuring tool to approximate the paved distance by the shadow. Therefore, the advancing speed of the shadow can be achieved by dividing the paved distance (the number of the lightened/darkened prisms multiplied by basal prism size) by time. Moreover, the prisms' size, packing, and geometry can roughly cancel the direction factor to make the estimated distance fairly direction-independent. Hence, regardless of direction, stationery, slow-moving and fast-moving objects can be differentiated and sorted by advancing speed.

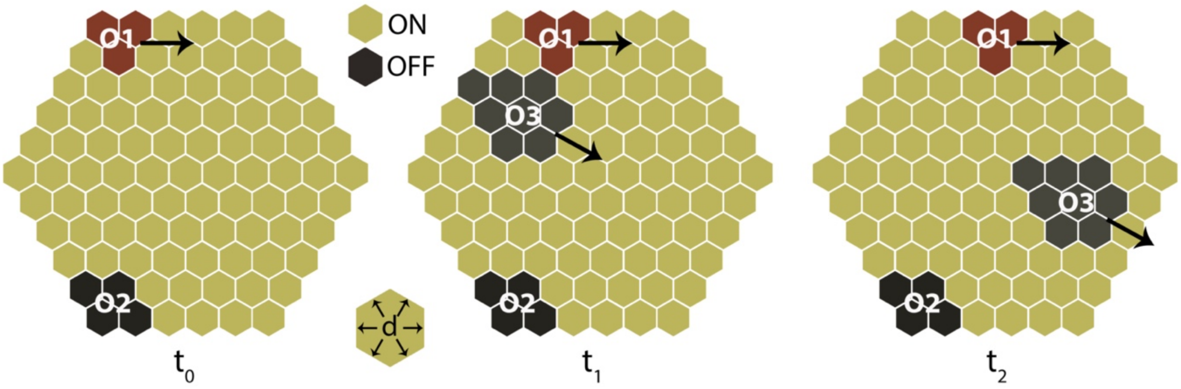


$$V_{O1} = \frac{2d}{t_2 - t_0}$$ Slow-moving object

$$V_{O2} = 0$$ Stationary object

$$V_{O3} = \frac{\sim 6d}{t_2 - t_1}$$ Fast-moving object
